# Spatially discrete signalling niches regulate fibroblast heterogeneity in human lung cancer

**DOI:** 10.1101/2020.06.08.134270

**Authors:** CJ Hanley, S Waise, R Parker, MA Lopez, J Taylor, LM Kimbley, J West, CH Ottensmeier, MJJ Rose-Zerilli, GJ Thomas

## Abstract

Fibroblasts are functionally heterogeneous cells, capable of promoting and suppressing tumour progression. Across cancer types, the extent and cause of this phenotypic diversity remains unknown. We used single-cell RNA sequencing and multiplexed immunohistochemistry to examine fibroblast heterogeneity in human lung and non-small cell lung cancer (NSCLC) samples. This identified seven fibroblast subpopulations: including inflammatory fibroblasts and myofibroblasts (representing terminal differentiation states), quiescent fibroblasts, proto-myofibroblasts (x2) and proto-inflammatory fibroblasts (x2). Fibroblast subpopulations were variably distributed throughout tissues but accumulated at discrete niches associated with differentiation status. Bioinformatics analyses suggested TGF-β1 and IL-1 as key regulators of myofibroblastic and inflammatory differentiation respectively. However, *in vitro* analyses showed that whilst TGF-β1 stimulation in combination with increased tissue tension could induce myofibroblast marker expression, it failed to fully recapitulate *ex-vivo* phenotypes. Similarly, IL-1β treatment only induced upregulation of a subset of inflammatory fibroblast marker genes. *In silico* modelling of ligand-receptor signalling identified additional pathways and cell interactions likely to be involved in fibroblast activation, which can be examined using publicly available R shiny applications (at the following links: **<u>myofibroblast activation</u>** and **<u>inflammatory fibroblast activation</u>**). This highlighted a potential role for IL-11 and IL-6 (among other ligands) in myofibroblast and inflammatory fibroblast activation respectively. This analysis provides valuable insight into fibroblast subtypes and differentiation mechanisms in NSCLC.

## Main text

Fibroblasts are abundant but poorly characterised cells that play pivotal roles in wound healing^1^, extracellular matrix (ECM) remodelling^2^ and inflammation^3^. In cancer, fibroblasts are the most common type of stromal cell and promote multiple hallmarks of malignancy^4-6^. These cells have been a putative therapeutic target in cancer for over a decade, but clinically effective strategies are yet to be identified. This is likely due to incomplete understanding of fibroblast heterogeneity and their precise role in tumour progression, which is exemplified by reports of both tumour-promoting and suppressive effects^7^. The diverse roles attributed to fibroblasts may be regulated by distinct subpopulations, such as those previously described in the dermis^8^, fibrotic lesions^9^ and in multiple solid cancers^10-12^. Cancer associated fibroblast (CAF) subpopulations are known to include myofibroblastic and inflammatory phenotypes^3, 11^. However, the extent of phenotypic diversity and the underlying molecular mechanisms regulating this heterogeneity remain unclear. Recent technical advances, such as single-cell RNA sequencing (scRNA-seq), have improved our understanding of the tumour microenvironment’s multicellular complexity^10, 13^, providing a platform to investigate unanswered questions in fibroblast and cancer biology.

We performed scRNA-seq on human lung tissue samples (n=18; 6 tumour-adjacent normal, 7 squamous cell carcinomas [LUSC], and 5 adenocarcinomas [LUAD]), using a previously-described protocol to enrich for fibroblasts^14^. Clustering and differential gene expression analyses identified distinct cell populations, representing mesenchymal, lymphocyte, myeloid and epithelial lineages (**Fig. 1a-b** and **Extended Data Fig. 1**). Within the mesenchymal lineage, we identified endothelial cells (marked by *VWF* and other canonical marker expression) and stromal cells (marked by expression of known fibroblast marker genes, *e.g. DCN, COL1A2* and *COL3A1*; **Fig. 1b** and **Extended Data Fig. 1e**).

**Figure 1:**
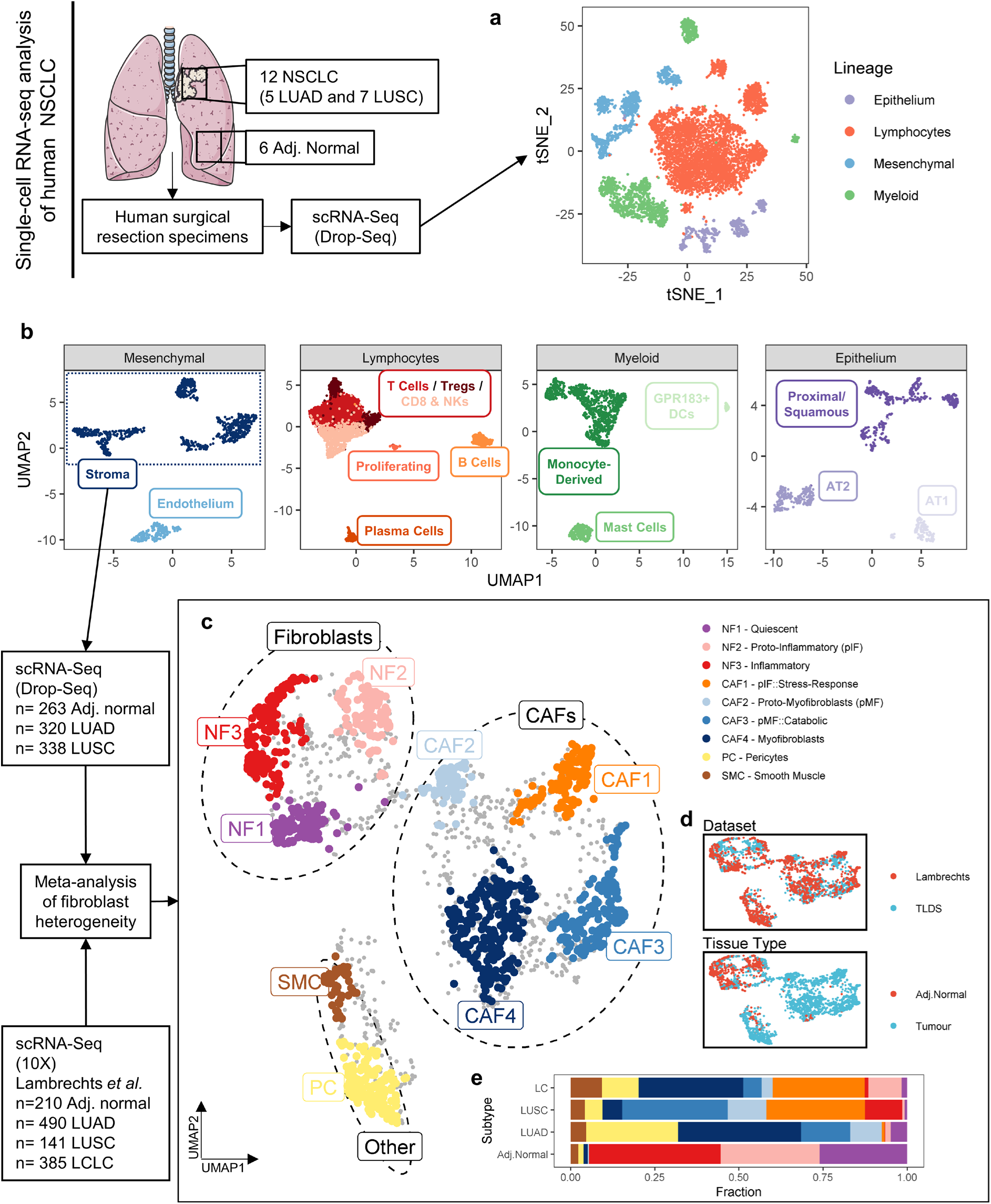
Single cell RNA-sequencing identifies fibroblast subpopulations in human NSCLC. **a)** Human tumour-adjacent normal lung (n=6) and non-small cell lung cancer (NSCLC) (n=12; adenocarcinoma [LUAD] or squamous cell carcinoma [LUSC]) samples were disaggregated and analysed by scRNA-seq. A 2D visualisation (tSNE dimensionality reduction) of cell-type clustering is shown, highlighting different cell lineages. Further analysis is shown in Extended Data Fig.1. **b)** UMAP plots showing subpopulations identified for each cell lineage identified in panel a. **c-e)** Transcriptome-based meta-analysis of mesenchymal stromal cells (predominantly fibroblasts) in human NSCLC (LUAD, LUSC or large cell lung cancer [LCLC]; further analysis shown in Extended Data Fig.2). **c)** A 2D visualisation (UMAP dimensionality reduction) showing unsupervised clustering of stromal cells, highlighting subpopulations (grey points represent non-clustered cells). **d)** UMAP plot (as per c) with points coloured by dataset or tissue type, as indicated. **e)** Stacked bar chart showing the relative abundance of each stromal subpopulation by NSCLC subtype.

To investigate fibroblast heterogeneity in NSCLC we performed a meta-analysis (see online methods), combining our stromal population (n=921) with transcriptomically similar cells from a publicly available NSCLC scRNA-seq data resource, generated by Lambrechts *et al*.^13^ (n=1226; total n=2147). Unsupervised clustering identified nine subpopulations (**Fig. 1c-e** and **Extended Data Fig. 2a-b**), representing three stromal cell groups: fibroblasts (NF1-3), predominantly isolated from tumour-adjacent normal tissue (74-77%); CAFs (CAF1-4), predominantly isolated from tumour samples (98-100%); and two non-fibroblastic populations (pericytes [PCs], identified by upregulation of *RGS5*^15^; and smooth muscle cells [SMCs], identified by *MYH11* upregulation; **Extended Data Fig. 2c**).

Many studies have attempted to identify a specific marker for CAFs, but this remains elusive. α-smooth muscle actin (coded for by *ACTA2*) is a commonly used CAF and myofibroblast marker^16^. However, when identifying myofibroblasts from scRNA-seq data, *ACTA2* is not a suitable (single) marker as it is highly expressed by pericytes and smooth muscle cells^17^. This has led to the misclassification of myofibroblasts in previous studies^10, 13, 18^. Instead, combined expression of *ACTA2* and ECM genes (*e.g. COL1A1*) is required for accurate myofibroblast classification from scRNA-seq data (**Extended Data Fig. 2c**).

To examine fibroblast phenotypes further we used multiple bioinformatics approaches. Cluster marker genes common to both datasets were identified (consensus markers; **Supplementary Table 1**) and examined for enrichment with genes from biological process gene ontology (GO) terms (**Fig. 2a** and **Supplementary Table 2**). Overlap between our scRNA-seq clusters and previously described fibroblast phenotypes was assessed by examining the differential expression of gene sets identified in previous studies (**Fig. 2b** and **Supplementary Table 3**). Diffusion map^19^ dimensionality reduction and the Slingshot^20^ algorithm was used to examine the continuum of consensus marker expression between fibroblasts and identify trajectory inferred cell-state transitions respectively (**Fig. 2c-d**). This enabled further identification (in addition to cluster markers) of genes associated with fibroblast heterogeneity, by modelling expression levels across multiple cells as a product of trajectory inferred “pseudotime”. Correlation network analysis of genes differentially expressed in pseudotime, along each trajectory, was then used to identify pseudotime gene expression modules (**Fig. 2e-f** and **Supplementary Tables 4&5**).

**Figure 2:**
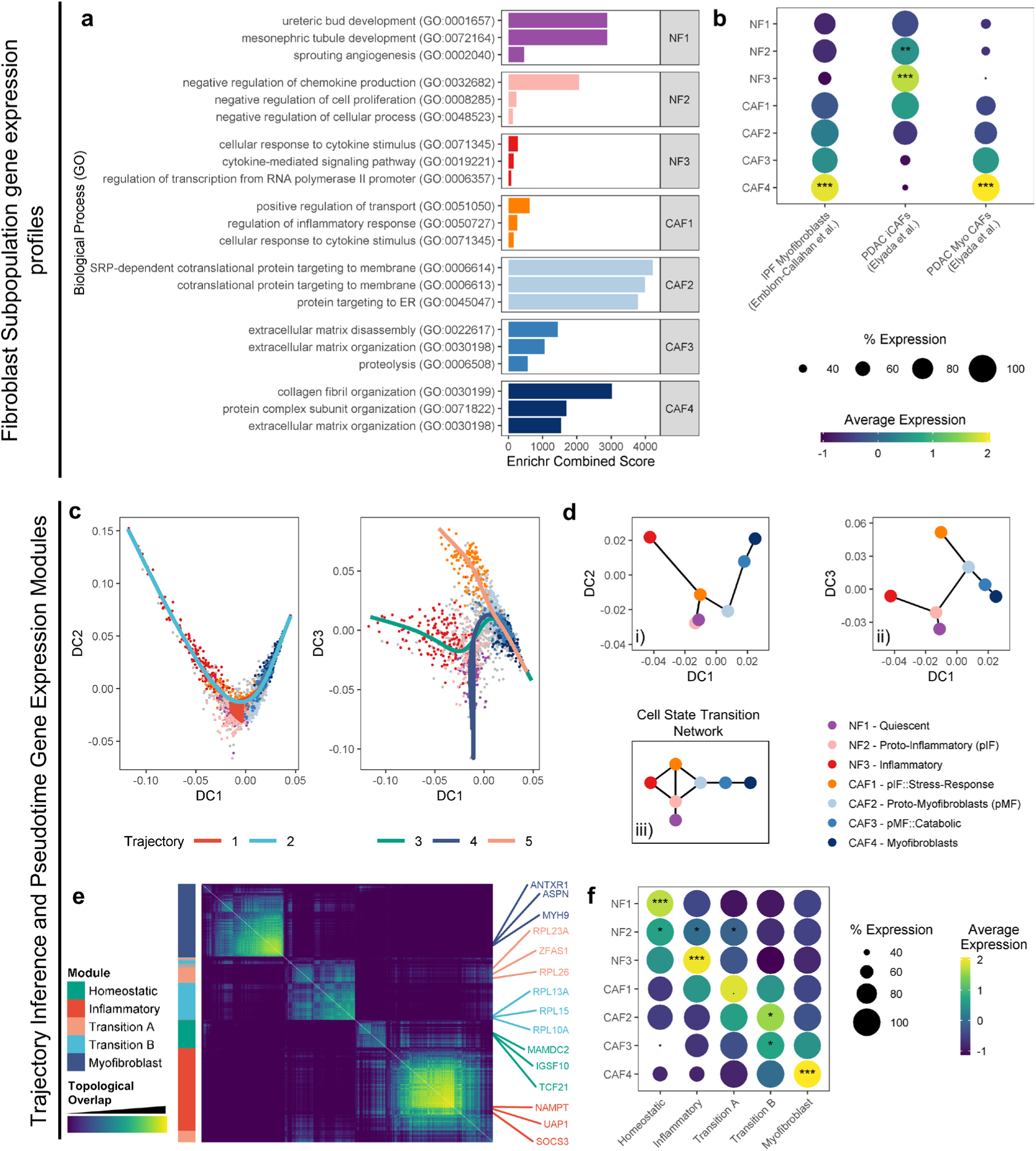
Fibroblast subpopulations represent a continuum of differentiation states ranging from inflammatory fibroblasts to myofibroblasts. **a)** Barplot showing the enrichr combined score for biological process (GO-terms) enrichment associated with consensus marker genes for each fibroblast subpopulation (complete results provided in Supplementary Tables 1&2). **b)** Dotplots showing fibroblast subpopulation’s expression of previously described fibroblast phenotype gene sets (complete results provided in Supplementary Table 3; Myo CAFs = Myofibroblastic-CAFs; iCAFs = inflammatory-CAFs; IPF = Idiopathic pulmonary fibrosis). Statistical significance was assessed using a Wilcox test with fdr correction to compare the median expression level per sample for each cluster to all other clusters (***adj. *p*<0.001, **adj. *p*<0.01, *adj. *p*<0.05). **c)** Diffusion map showing trajectory inference using the slingshot algorithm applied to components 1 and 2 or 1 and 3. Points represent individual cells coloured by cluster and lines represent the principal curves identified for each inferred trajectory. **d)** Network plots summarising all cell state transitions identified by trajectory inference. Nodes represent cluster medians mapped to diffusion map components (as shown in c) 1 and 2 (i); 1 and 3 (ii); or in a tree layout (iii) showing all cell state transitions identified. **e)** Heatmap showing topological overlap of gene expression in trajectory-inferred pseudotime as a symmetrical matrix. Gene modules, identified by correlation network analysis, are shown to the left of the heatmap. The top 3 hub genes (measured by intra-module connectivity) for each module are labelled. **f)** Dotplots showing the expression of each pseudotime gene expression module by different fibroblast subpopulations. Statistical significance was assessed using a Wilcox test with fdr correction to compare the median expression level per sample for each cluster to all other clusters (***adj. *p*<0.001, **adj. *p*<0.01, *adj. *p*<0.05, ^•^ adj. *p*=0.05).

Similar to scRNA-seq analysis of human head and neck squamous cell carcinoma^10^ and pancreatic adenocarcinoma^11, 12^ we identified a myofibroblastic-CAF subpopulation (CAF4) with upregulated expression of previously described CAF markers (*e.g. ACTA2, POSTN* and *FAP*). Multiple studies have documented common features between fibroblast activation in cancer and during wound healing or fibrosis^21^. Consistent with this, we found that myofibroblastic-CAF significantly upregulated a gene signature associated with end-stage idiopathic pulmonary fibrosis^22^ (IPF; **Fig. 2b** and **Supplementary Table 3**).

Trajectory inference identified two intermediate states in myofibroblast differentiation (CAF2 and CAF3; **Fig. 2c-d**). These subpopulations upregulated previously described CAF markers (*e.g. CXCL12, IGF1* and *COL1A1* -CAF2; *POSTN, FAP* and *COL1A1* -CAF3; **Supplementary Table 1**), without increased *ACTA2* expression, suggesting they represent “proto-myofibroblast” phenotypes^23^ (**Extended Data Fig. 2c** and **Supplementary Table 1**). The dynamic transition to a fully differentiated myofibroblast phenotype was captured by a pseudotime gene expression module (“Transition B”), which was significantly upregulated by both proto-myofibroblast populations (CAF2 and CAF3; **Fig. 2e-f**). This module and the CAF2 markers included multiple ribosomal genes (**Fig. 2a** and **Supplementary Tables 1&5**), which may reflect development of rough endoplasmic reticulum: a prominent ultrastructural feature of myofibroblasts^24^. In addition, CAF3 markers were enriched for genes involved in catabolic processes, including multiple matrix metalloproteases (MMPs; *e.g. MMP1, MMP11, MMP3, MMP14*; **Fig. 2a** and **Extended Data Fig. 2d**). CAF3 may therefore represent a novel “catabolic” CAF subpopulation with matrix remodelling functions. Pseudotime gene expression analysis identified genes increasingly expressed as cells differentiate from CAF2-CAF3-CAF4 (**Fig. 2d-f**). This “Myofibroblast” module was enriched for genes upregulated by fibroblasts treated with TGF-β; genes involved in ECM-receptor interactions and focal adhesion pathways; and multiple targets of the miR-29 family of micro RNAs (**Fig. 2e-f**; **Supplementary Table 5**).

Inflammatory CAFs (iCAFs) have been shown to represent a subpopulation distinct from myofibroblasts in pancreatic cancer^11, 12^. We identified a similar inflammatory subpopulation, predominantly found in tumour-adjacent normal tissues (NF3; **Fig. 2b**). Trajectory inference showed inflammatory fibroblasts and myofibroblasts represent terminal differentiation states in a fibroblast phenotype continuum (**Fig. 2c&d**).

Similar to myofibroblast differentiation, trajectory inference identified two subpopulations as likely precursors to inflammatory fibroblasts (NF2 and CAF1; **Fig. 2c-d**). These were termed “proto-inflammatory fibroblasts”, for consistency with proto-myofibroblasts. Both proto-inflammatory populations up-regulated previously described iCAF marker genes (*e.g. IGFBP6, MFAP5 and GSN* -NF2; *SGK1, APOE* and *CXCL2* -CAF1; *C3* and *PLA2G2A* -NF2 and CAF1). As these proto-inflammatory populations were largely restricted to either tumour-adjacent normal samples (NF2) or tumour samples (CAF1) it is possible that distinct processes can regulate inflammatory phenotypes in fibroblasts. Notably, the CAF1 subpopulation also upregulated genes involved in stress response signalling (*e.g. HIF1A, MT1X* and *MT1E*), consistent with previous studies showing that cellular stress can induce an inflammatory fibroblast phenotype^25, 26^. Pseudotime gene expression analysis identified genes increasingly expressed as cells differentiate into inflammatory fibroblasts. This “inflammatory” module was enriched for genes upregulated by fibroblasts treated with IL-1β; genes involved in TNF and IL-17 signalling pathways; transcription factors RELA and STAT3; and miR-98-5p (**Supplementary Table 5**).

To validate scRNA-seq findings we examined the spatial distribution of stromal subpopulations within the NSCLC microenvironment, using histo-cytometry analysis of multiplexed immunohistochemistry whole slide imaging (**Fig. 3a**). Pan-cytokeratin was used as an epithelial (and stromal cell exclusion) marker. ACTA2, POSTN and SERPINE1 were selected as stromal cell markers, based on their variable expression between stromal subpopulations (**Extended Data Fig. 3a**).

**Figure 3:**
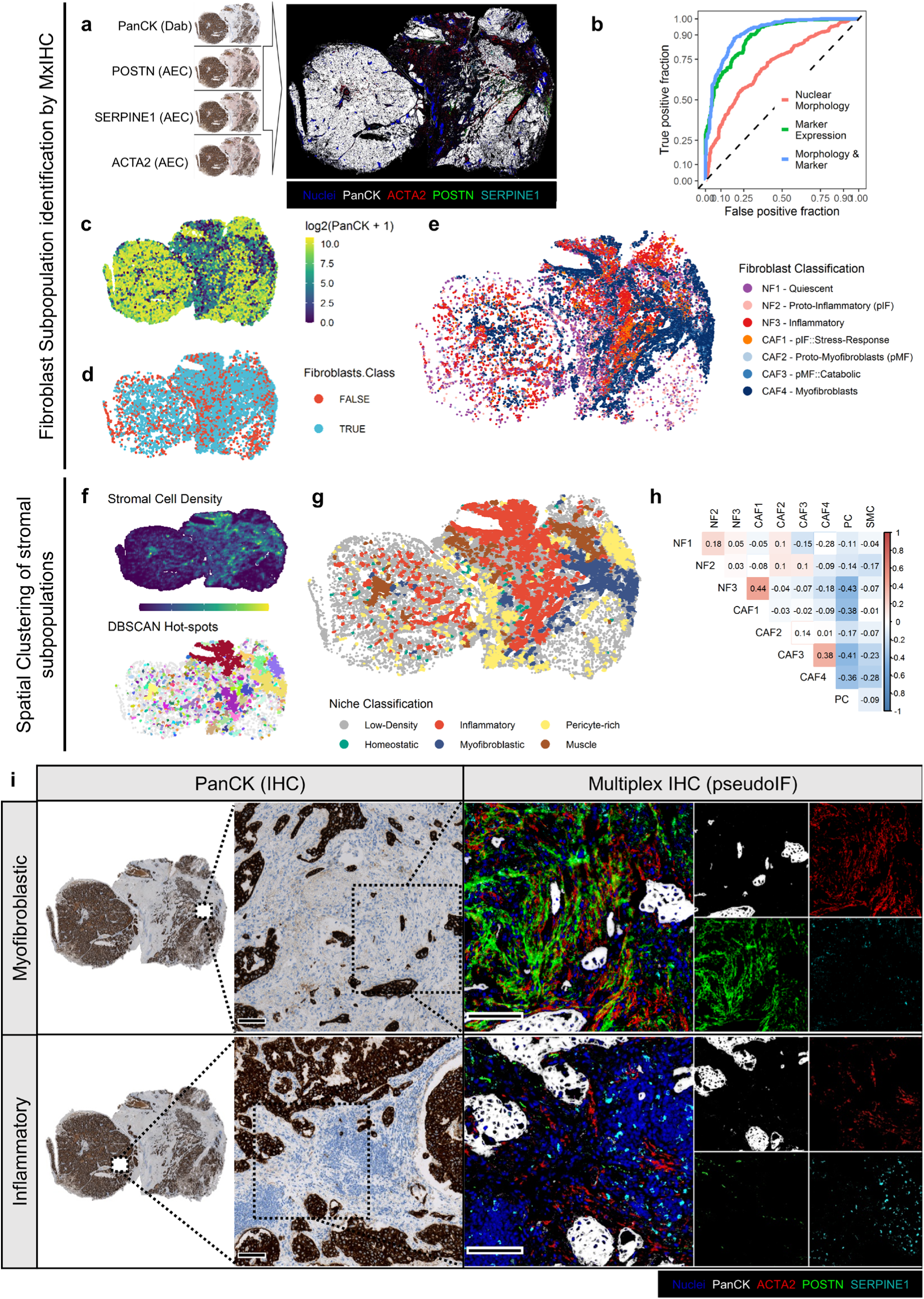
Fibroblast subpopulations accumulate at spatially discrete niches. **a)** Representative example of multiplex immunohistochemistry (mxIHC) staining and pseudo-immunofluorescence (pIF) image generation. **b)** ROC analysis of fibroblast detection classifiers trained using the metrics indicated (see also Extended Data Fig. 3b). **c)** Representative point pattern plot showing per cell expression levels of pan-cytokeratin (PanCK). Sample dataset was randomly downsampled to 10% of cells for plotting. **d)** As per c showing results of the fibroblast classifier. Only non-epithelial cells (PanCK low) are shown; the sample dataset was randomly downsampled to 10% for plotting. **e)** As per c showing fibroblast subpopulation classifier results. **f)** Representative images showing stromal hot-spot identification. The upper panel shows a heatmap for stromal cell density; the lower panel shows the results from DBSCAN clustering analysis. **g)** Representative point pattern plot showing the results of Niche classification by clustering stromal hotspots based on subpopulation composition (see also Extended Data Fig.3c) **h)** Heatmap showing the pairwise spearman correlation between stromal subpopulations across DBSCAN hotspots. Non-significant (adj. *p*>0.05) correlations are shown in white squares. **i)** Representative micrographs showing myofibroblastic and inflammatory niches. Scale bars represent 100 µm.

To detect stromal cells, we developed a machine learning (random forest) classifier (see online methods). Optimisation showed this classifier could detect manually annotated stromal cells based on marker expression and nuclear morphology with 84.4% accuracy (area under the ROC curve = 0.92; **Fig. 3b-d** and **Extended Data Fig. 3b**). This identified 1.9×10^6^ stromal cells in the histo-cytometry dataset, which were then classified into the relevant subpopulations using a classifier trained on scRNA-seq data (**Fig. 3e** and **Extended Data Fig. 3a**).

To examine stromal cell distribution throughout whole slide images, the DBSCAN^27^ algorithm was used to identify spatially discrete regions of high stromal cell density (“hotspots”) in each tissue section (**Fig. 3f**). Clustering of stromal cell hotspots showed that fibroblast activation status was associated with spatially discrete niches, consistent with the scRNA-seq pseudotime gene expression modules (**Fig. 3g-h** and **Extended Data Fig. 3c**). This identified three types of fibroblast niche: homeostatic, where quiescent (NF1), proto-inflammatory (NF2) or proto-myofibroblasts (CAF2) were found; inflammatory, with high levels of inflammatory (NF3) and stress-response proto-inflammatory (CAF1) fibroblasts; and myofibroblastic, with increased accumulation of myofibroblasts (CAF4) and catabolic proto-myofibroblasts (CAF3; **Fig. 3i** and **Extended Data Fig. 3c**). Regions with high levels of pericyte/pericyte-derived CAFs or SMCs were typically associated with the presence of large vessels (**Extended Data Fig. 3d**).

As inflammatory and myofibroblastic CAFs have previously described roles in tumour progression^3, 21^ and represent terminal differentiation states, we sought to identify the molecular mechanisms responsible for their divergent activation. NicheNet^28^ and enrichr^29^ were used to identify ligands previously described to regulate the expression of markers or pseudotime module genes for myofibroblasts or inflammatory fibroblasts. Consistent with previous studies^21, 30^, this highlighted TGF-β1 and IL-1β as likely regulators of myofibroblast and inflammatory phenotypes respectively (**Fig.4 a-b** and **Extended Data Fig. 4a-b**). In our scRNA-seq data *TGFB1* was variably expressed by different cell types; whereas *IL1B* was predominantly expressed by myeloid cells suggesting a potential paracrine signalling mechanism (**Extended Data Fig. 4c**). To test the effect of these cytokines on fibroblast differentiation, primary human fibroblasts isolated from tumour-adjacent lung tissue were cultured on collagen-coated plates with physiological lung tissue stiffness (2 kPa elastic modulus^31^) and treated with IL-1β (10 ng/ml for 72 hours) or TGF-β1 (2 ng/ml for 72 hours), reproducing conditions used for manipulating fibroblast phenotypes in previous studies^21, 32^.

As expected, TGF-β1 treatment significantly increased expression of markers from all subpopulations on the myofibroblast activation trajectory (CAF2, CAF3 and CAF4; **Fig. 4c**). To determine whether these changes in gene expression induced *in vitro* were physiologically relevant, we compared marker gene expression levels to *ex vivo* phenotypes (**Fig. 4d-e**). *In vitro* control fibroblasts were found to most closely resemble proto-myofibroblasts (CAF2; **Fig. 4e**). TGF-β1 treatment had minimal impact on the differentiation status of these cells, when compared to *ex vivo* fibroblasts (**Fig. 4d-e**), with only 26% of myofibroblast (CAF4) marker genes and 17% of the myofibroblast pseudotime module significantly upregulated (**Extended Data Fig. 4f** and **Supplementary Table 6**).

**Figure 4:**
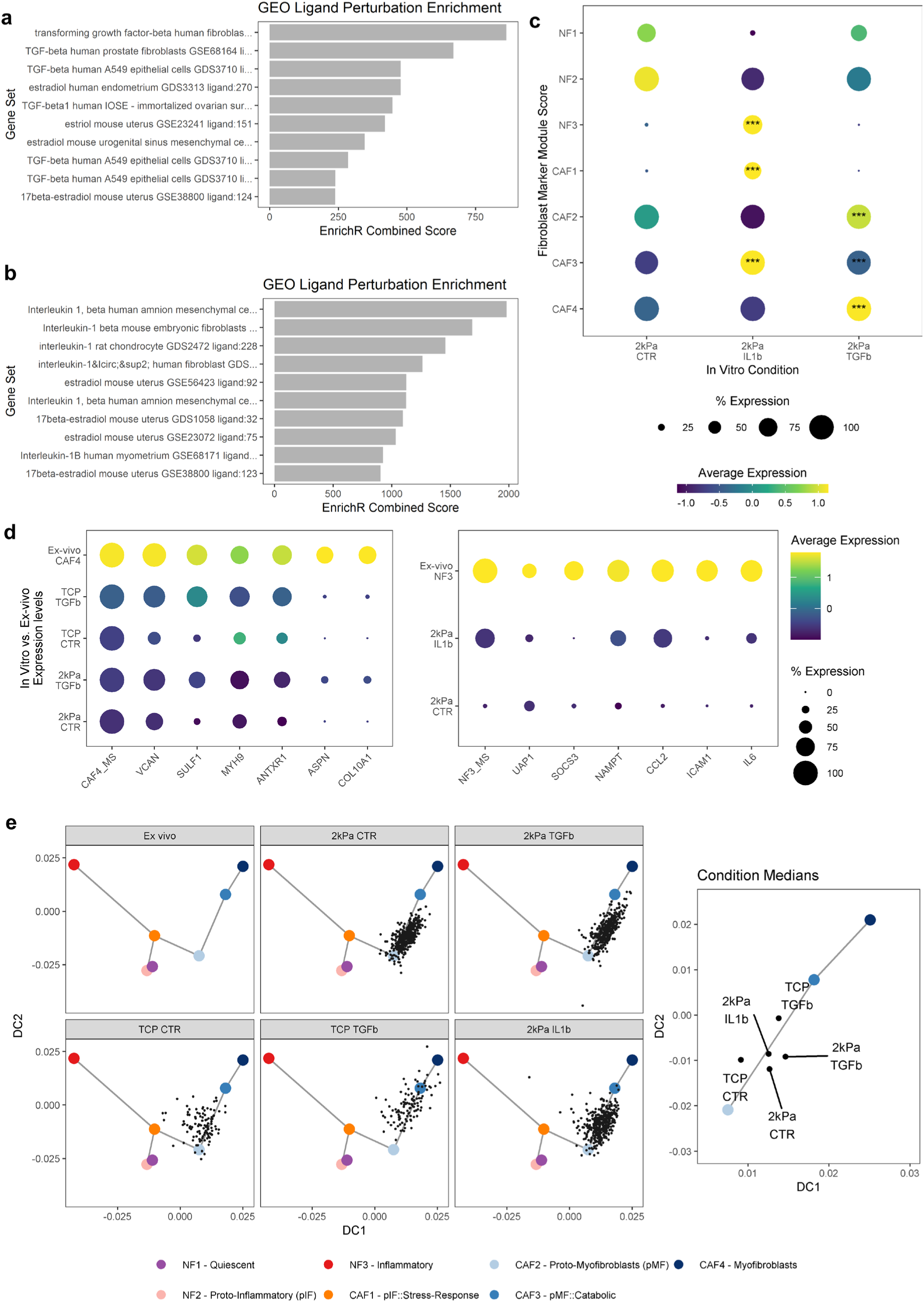
IL-1β and TGF-β1 treatment alone is not sufficient to generate ex-vivo inflammatory and myofibroblast phenotypes respectively. **a-b)** Predicted ligand involvement in Myofibroblast (a) and inflammatory fibroblast (b) gene expression. Bar plots show enrichment scores for “GEO ligand perturbation UP” gene sets from the enrichr database. **c)** Dotplots showing expression of fibroblast subpopulation marker genes, measured by scRNA-seq, in primary fibroblasts cultured *in vitro* on a “soft” (2kPa) substrate and treated with IL-1β (10ng/ml) or TGF-β1 (2ng/ml). statistical significance was assessed using a wilcox test compared to the control (CTR) group with fdr correction (****adj. *p*<0.001, ***adj. *p*<0.001). **d)** Dotplots comparing *ex vivo* and *in vitro* expression levels for myofibroblast (CAF4) or inflammatory fibroblast (NF3) marker genes. Fibroblasts were treated as per panel c or cultured on tissue culture plastic (TCP). **e)** Comparison of *in vitro* and *ex vivo* fibroblast phenotypes by projection onto the fibroblast marker gene diffusion map (shown in Fig.2d). Points represent individual cells or the median position of cells from each condition.

Tissue tension impacts myofibroblast differentiation in response to TGF-β1^33^ and the myofibroblast pseudotime module was enriched for pathways involved in cell-substrate interactions (**Supplementary Table 5**). Therefore, we hypothesised that increasing substrate stiffness would enhance the effect of TGF-β1 treatment. This was confirmed by comparing primary fibroblasts grown on either a substrate with physiological stiffness (2 kPa) or tissue culture plastic (TCP, ∼2 GPa). Increased substrate stiffness alone induced comparable upregulation of myofibroblast marker genes to TGF-β1 treatment (**Extended Data Fig. 4d-e**). The combination of TCP and TGF-β1 treatment caused the largest increase in myofibroblast gene expression. However, this still failed to fully induce the level of myofibroblast differentiation observed *ex-vivo* (upregulating 39% of CAF4 marker genes and 29% of the myofibroblast pseudotime module; **Fig. 4d-e**, Extended Data Fig. 4f and **Supplementary Table 6**).

Similar to TGF-β1 treatment, IL-1β caused a significant increase in the expression of a subset of genes up-regulated by *ex vivo* inflammatory fibroblasts (upregulating 14% of NF3 markers and 13% of the inflammatory pseudotime module; **Fig. 4c-e** and **Extended Data Fig. 4g**). Notably, IL-1β also significantly increased the expression of stress-response (CAF1) and catabolic (CAF3) marker genes (28% and 47% respectively; **Fig. 4c**).

To align these findings with previous studies, we curated lists of IL-1β or TGF-β1 dependent inflammatory fibroblast or myofibroblast genes by combining our findings with publicly available data (**Supplementary Table 6**). This showed that, across multiple studies, TGF-β1 has been found to upregulate 59% of myofibroblast genes (markers and pseudotime module) and IL-1β 32% of Inflammatory fibroblast genes (**Extended Data Fig. 4f-g**). Together, these data indicate myofibroblast and inflammatory fibroblast activation in NSCLC is only partially TGF-β1- or IL-1β-dependent.

To identify additional stimuli regulating fibroblast differentiation *in vivo*, we analysed the expression of ligand-receptor pairs in our scRNA-seq dataset. To compare fibroblast phenotypes between samples, the median position of fibroblasts in the diffusion map dimensionality reduction was used (**Fig. 5a**). We then examined which cells were responsible for expressing ligands recognised by fibroblast receptors. This showed that in myofibroblast or proto-myofibroblast rich samples, fibroblasts themselves were the primary ligand source - suggesting autocrine signalling is likely critical to regulating these phenotypes (**Fig. 5b**). In contrast, epithelial, myeloid cells and fibroblasts all contributed similarly to ligand expression in inflammatory or proto-inflammatory rich samples (**Fig. 5b**).

**Figure 5:**
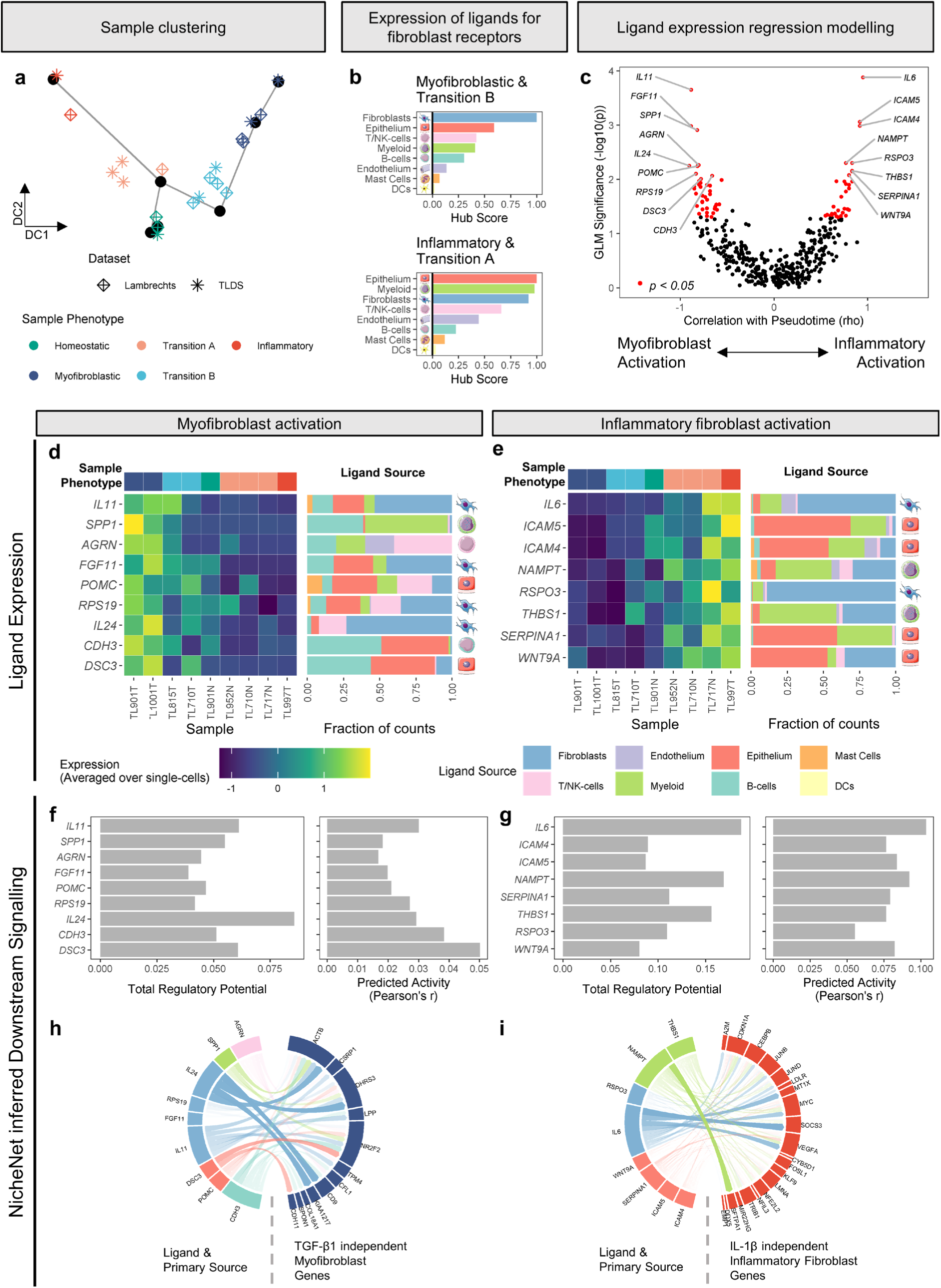
In Silico *modelling identifies IL-1β and TGF-β1 independent mechanisms of fibroblast activation*. **a)** Diffusion map showing sample clustering based on fibroblast phenotypes. Samples are represented by the median position of fibroblasts in the diffusion map. **b)** Bar plots show the Kleinberg Hub Scores for expression of ligands recognised by fibroblast expressed receptors in different sample types. **c)** Volcano plot showing the results from linear regression modelling of associations between ligand expression (per sample averaged over single cells) and each sample’s median fibroblast phenotype. Ligands with *p*<0.01 are labelled, **d-e)** Heatmap showing the average expression of ligands associated with myofibroblasts (d) or inflammatory fibroblast (e) activation and barplots showing the relative contribution of different cell types to this expression. **f)** Barplots showing the regulatory potential of ligands associated with myofibroblast activation on TGF-β1 independent myofibroblast genes. **g)** Barplots showing the regulatory potential of ligands associated with inflammatory fibroblast activation on Il-1β independent inflammatory genes. **h)** Circos plot showing known links between ligands associated with myofibroblast activation and TGF-β1 independent myofibroblast genes. The primary source of each ligand is also shown. **i)** Circos plot showing known links between ligands associated with inflammatory fibroblast activation and Il-1β independent inflammatory genes. The primary source of each ligand is also shown.

To examine the potential role of specific ligands in fibroblast activation, we tested the predictive value of each ligand’s average expression (per sample) on the sample’s position in pseudotime, using generalised linear modelling (GLM; **Fig. 5c**). Ligands identified as significant (*p*<0.05) in this modelling were then examined for previously reported regulatory potential over IL-1β- or TGF-β1-independent inflammatory or myofibroblast genes respectively, using the NicheNet database^28^ (**Supplementary Table 8**; further analysis of ligands with *p<0.01* is shown in **Fig. 5c-i**). This identified multiple ligands with potential to regulate TGF-β1- or IL-1β-independent features of myofibroblast and inflammatory fibroblast activation respectively (including IL11, SPP1, AGRN – myofibroblast; IL6, ICAM5, ICAM4 – inflammatory; **Fig. 5c-i**). The results from this analysis for all ligands with *p<*0.05, including inferred downstream signalling pathways, can be examined interactively using R shiny apps available at the following links: <u>myofibroblast activation</u> and <u>inflammatory fibroblast activation</u> (examples shown in **Extended Data Fig. 5**).

In summary, we have characterised fibroblast heterogeneity in human NSCLC and tumour-adjacent tissues. We identified seven subpopulations, with varying abundance between NSCLC subtypes. These were shown to accumulate at spatially discrete signalling niches, within tumours, which may be associated with functional synergy. We also elucidated mechanisms regulating myofibroblast and inflammatory fibroblast activation in the NSCLC tumour microenvironment, by combining *in vitro* and *in silico* modelling. Fully differentiated myofibroblasts, which are well described to impact patient survival across solid cancers^21^, were more common in LUAD than LUSC. Notably, we found myofibroblasts and catabolic proto-myofibroblasts occupy similar spatial niches and therefore may play complimentary roles in ECM deposition and degradation. Consequently, their relative contribution to a niche may differentially impact tumour cell invasion^34^. Our analysis suggests TGF-β1 triggers myofibroblast differentiation, but additional stimuli are required to complete the process. This is consistent with previous studies showing that TGF-β1 is not required to maintain myofibroblast activation^5^. The ligands we identified as associated with myofibroblast activation are likely to represent these additional stimuli (**Fig. 5d, f&h**). For example, IL-11 inhibition can revert myofibroblast phenotypes in murine models of IPF^35^. Inflammatory fibroblasts are also a prominent feature of the NSCLC tumour microenvironment. In addition to the previously described role for IL-1 in inflammatory fibroblast activation^30^, we showed that IL-1β induced expression of markers for all subpopulations predominantly isolated from LUSC samples (NF3, CAF1 and CAF3). LUSC have higher levels of necrosis and macrophage infiltration than LUAD^36^. Consistent with this as we found myeloid cells to be the primary source of *IL1B*, suggesting paracrine signalling interactions may play an important role in regulating fibroblast heterogeneity between NSCLC subtypes. However, similar to myofibroblast differentiation, complete inflammatory fibroblast activation requires stimuli in addition to IL-1β. Notably, we identified a potential role for IL6 in this process (**Fig. 5e, g&i**) and showed that *IL6* expression is increased in fibroblasts following IL-1β treatment (**Extended Data Fig. 4d**). Therefore, it is possible that autocrine IL-6 signalling fully activates inflammatory fibroblasts, following initial stimulation by myeloid-cell derived IL-1β (**Extended Data Fig. 4c**). These findings provide novel insights into the signalling mechanisms that regulate fibroblast heterogeneity in human NSCLC. These are important considerations for future functional analyses and may allow more precise therapeutic strategies for fibroblast targeting to be developed.

## Acknowledgements

This work was supported by Cancer Research UK (CRUK) and Medical Research Council (MRC) Clinical Research Training Fellowships and a Pathological Society Trainee’s Small Grant (S.W.); a CRUK accelerator award (A20256; G.J.T.); a British Lung Foundation pump-priming research grant (PPRG17-12; C.J.H); an MRC Discovery award (MC_PC_15078; M.J.J.R-Z and J.W.); a Leuka Charity John Goldman Fellowship for Future Science (2016/JGF/0003; M.J.J.R-Z.); and a Southampton Cancer Research UK Centre Development Fund Award (M.J.J.R-Z., C.H.O., J.W., C.J.H. & G.J.T.). The authors thank Evan Macosko, Melissa Goldman and Steve McCarroll, for their helpful advice implementing Drop-Seq; Dr Martin Fischlechner for his work developing the microfluidic hardware; Dr. Serena Chee (University Hospital Southampton), Benjamin Johnson, Carine Fixmer and Maria Lane (TargetLung Clinical Trials Associates) for enabling access to clinical samples; and the patients involved in this study.

## Author Contributions

C.J.H. and G.J.T. conceived and supervised the study; C.J.H., M.J.J.R-Z. and G.J.T. designed experiments; J.W. provided the microfluidic hardware; C.J.H., S.W., R.P., J.T. and M.A.L. performed experiments; C.H.O. provided access to clinical samples; C.J.H., G.J.T., M.J.J.R-Z. and S.W. analysed data; C.J.H., G.J.T. and S.W. wrote the manuscript; C.H.O. and M.J.J.R-Z. made manuscript revisions.

## Methods

### Sample acquisition and processing

Lung samples were received from the TargetLung study (REC number 14/SC/0186; Table S8) as described previously^14^ and transported (within 1 hour) to the laboratory on ice in serum-free Dulbecco’s Modified Eagle Medium (DMEM; Sigma-Aldrich).

Tissue disaggregation was performed as previously described^14^. Briefly, samples were washed, incised and incubated with Collagenase P (3 U/ml; Sigma) at 37 °C with agitation (200 rpm) for 60 minutes. The resulting suspension was strained; incubated with red cell lysis buffer (BioLegend); and re-suspended in PBS supplemented with 9% Optiprep (Sigma) and 0.1% bovine serum albumin (BSA). Single-cell transcriptome encapsulation was performed using a custom microfluidic platform (Drop-seq) as described previously^14, 37^.

### Single-cell RNA-seq data processing and analysis

#### “Target Lung” Dataset (TLDS) processing all cells

ScRNA-seq data processing and analysis was performed using the Seurat package in R (version 2.3.4)^38^, unless otherwise stated. Initial quality control was carried out to remove low quality events. First, we used a random forest classifier to exclude empty droplets as described previously^14, 39^. We then identified outliers for the fraction of reads mapping to mitochondrial genes (> 2 median absolute deviations; MADs) to exclude apoptotic cells.

Initial clustering was performed using a subset of genes selected based on variance and average non-zero expression excluding extreme outliers. Raw counts data was transformed (log((counts+1)/10000)) and scaled (regressing out nUMI) before performing PCA. Clusters were identified with the *FindClusters* function, using principal components identified as significant with *JackStraw* analysis (*p<*1e-5) and a resolution of 0.2. Cluster markers were identified using the *FindAllMarkers* function (ROC classifier) and cell types were assigned based on canonical marker expression or by significant enrichment (adj. *p*<0.0001) in previously described cell type markers from the Immunological Genome^40^ and LungGENS^41^ projects, assessed using the ToppFun gene set enrichment tool^42^.

This clustering identified a large cluster of cells comprised of lymphocytes. However, due to the relatively low nGene associated with lymphocytes (described previously^13^) and therefore high susceptibility to false negatives in marker detection (due to drop-out), this cluster had very few genes identified as markers. Therefore, it was unclear whether the lymphocytes were separated from low-quality droplets. To identify lymphocytes within this cluster the AddModuleScore function was used to calculate the average expression of previously described T-cell markers (*TRBC2, CD3D, CD3E, CD3G, CD2, IL7R, CD8A*) and NK cell markers (*FGFBP2, SPON2, KLRF1, NKG7, PRF1, KLRD1*). This cluster was then filtered further to remove cells negative for both gene signatures. This filtered dataset was then re-clustered as described above.

#### “Lambrechts” Dataset processing all cells

The “Lambrechts” scRNA-seq dataset^13^ was downloaded as the “all cells” .loom file from https://gbiomed.kuleuven.be/scRNAseq-NSCLC. The empty droplet classifier (described above) was applied to this data set, filtering out 4768 cells, leaving a filtered dataset consisting of 47930 cells. The *BuildRFClassifier* function was then used to build a random forest classifier for detecting fibroblasts in the Target Lung dataset from the variable genes used for cell clustering. This classifier was then used to detect fibroblasts within the Thienpont dataset for further analysis (described below).

#### Fibroblast clustering Meta-analysis

Variable genes were identified as described above for each fibroblast dataset (Target Lung and Thienpont). The genes identified as variable in both datasets were then combined using canonical correlation analysis (CCA) implemented in Seurat. Clustering was performed using canonical vectors (selected based on shared correlation values >0.1 using the *MetageneBicorPlot* function) and a resolution of 1. Consensus cluster markers were identified using the *FindAllMarkers* function (as described below). Initial clustering yielded a small cluster likely to represent cells that had been incorrectly clustered/classified as they were marked by expression of immune cell markers (*e.g. IL7R, CD3D, LYZ* and *PTPRC*). This cluster was therefore excluded, and the remaining cells were then re-clustered as described above using consensus cluster and node markers for CCA.

We then examined the connectivity, within the SNN graph used for clustering, between cells assigned to each cluster. Mixture model analysis (mixtools package in R^43^) of intra-cluster connectivity values was then performed, identifying a bimodal distribution representing highly connected “Hub cells” and less connected cells (likely between differentiation states). Cells with a probability <0.75 of association with the second (more connected) component were removed from the respective cluster.

#### Fibroblast Subpopulation Consensus Marker identification

Marker genes were identified for each dataset separately, using the *FindAllMarkers* function (MAST test, with default settings treating nUMI and ambient RNA content as latent variables). “Meta” logFC and adj. *p* values were then calculated as the minimum absolute log(fold change) or maximum adj. *p* value across the two datasets and genes were considered consensus markers where the meta-adj. *p* < 0.05 and meta-logFC > 0.25.

#### Trajectory inference

Consensus Marker genes were used for trajectory inference. To mitigate batch effects and technical confounders the dataset (TLDS or Lambrechts) and nUMI were “regressed out” using the ScaleData function in Seurat. Diffusion map dimensionality reduction was then performed on the scaled data, using the destiny R package. Trajectory inference and pseudotime ordering was carried out using the Slingshot package in R^20^. Inferred lineages were variable, dependent on the diffusion map components analysed. Therefore, we considered all lineages identified from analysing the first three diffusion map components as potential cell-state transitions (**Fig.2 c-d**).

To validate these findings we also performed trajectory inference using the DDRtree dimensionality reduction and minimum spanning tree cell ordering algorithm implemented in the Monocle R package (v2.8.0)^44^. This identified lineages consistent with those identified from analysis of the first two components in the diffusion map approach (**Extended Data Fig.2e**).

Pseudotime gene expression analysis was performed for each of the trajectories identified. Each dataset was tested separately, using a generalised additive model (GAM; Loess) to determine the significance of pseudotime as an explanatory variable for each gene’s expression. The maximum (least significant) value from analysing the two datasets was used a as a meta-*p* value, which was then adjusted to account for multiple hypothesis testing using the FDR correction.

Genes differentially expressed in pseudotime (meta-adj. *p* < 0.05) were examined further to identify pseudotime gene expression modules. Fitted values from the GAM model generated for each gene and each trajectory were used to perform gene correlation network analysis, using the WGCNA package in R^45^. The topological overlap (TO) between genes was calculated from a signed-hybrid adjacency matrix. Gene modules were identified using a dynamic tree cutting algorithm applied to a 1-TO distance matrix. Hub Genes were identified as those with greatest intra-modular connectivity.

#### Gene set enrichment analysis

The enrichr R package^29^ was used for assessing consensus marker genes and pseudotime gene expression modules for gene set enrichment.

#### Ligand-receptor interaction analysis

Genes with potential activity as receptor ligands were identified using the NicheNet database^28^. Receptors were considered to be expressed by fibroblasts if they were detected in at least 5% of fibroblasts (as measured by scRNA-seq). Sample phenotypes (either position in diffusion map 3-dimensional space or trajectory inferred pseudotime) were calculated using the median value for all fibroblasts isolated from each sample (samples with less than 20 fibroblasts were excluded from this analysis). Cell-type specific contributions to ligand expression were calculated as the proportion of counts for each sample (or sample group) detected from individual cell types. Average expression (AE) values for ligands per sample were calculated using the *AverageExpression* function in Seurat. These values were then transformed to log(AE+1) and used as predictor variables for trajectory inferred pseudotime in GLM modelling.

Signalling pathways were inferred for each ligand’s potential role in target gene regulation using the NicheNet database^28^ and enrichr gene set enrichment analyses. First, transcription factor databases (“ENCODE_TF_ChIP-seq_2015” and “ARCHS4_TFs_Coexp”) were examined for enrichment in target genes (myofibroblast or inflammatory genes) using enrichr^29^. Transcription factors with significant (adj.*p* < 0.05) enrichment in either of these databases were classed as “Target enriched TFs” (TE-TFs). To minimise overlap in pathway inference between myofibroblast and inflammatory fibroblast activation, where TE-TFs were significant for both sets of target genes the odds ratio was used to determine which pathway it was most likely associated with. The NicheNet ligand_tf_matrix was then used to determine whether these TE-TFs act downstream of the ligand under investigation (“active TE-TFs”). To simplify pathway inference the active TE-TFs were restricted to the top 5, ranked by their score in the ligand_tf_matrix. The NicheNet weighted ligand-signaling network was then filtered to only contain the ligand under investigation and genes detected in at least 5% of fibroblasts (by scRNA-seq). The shortest possible paths from ligand to active TE-TFs in this filtered network were then calculated using the *shortest_paths* function in the igraph R package^46^ (weighted by 1/weight).

### Multiplex Immunohistochemistry (MxIHC)

#### Staining

Immunohistochemical staining of sections from ten TargetLung patients was performed, using a previously-described multiplexed protocol^47^. Four micrometre sections of formalin-fixed paraffin-embedded sections were mounted on Superfrost slides (ThermoFisher) and baked for 60 minutes at 60 °C. Deparaffinisation, rehydration, antigen retrieval and imunihistochemical staining were performed using the PT Link Autostainer (Dako) pre-defined program. Antigen retrieval for all antibodies was performed using the EnVision FLEX Target Retrieval Solution, High pH (Dako).

Sections were incubated with primary antibody (anti-pan-cytokeratin, clone AE1/AE3, Dako, pre-diluted; anti-POSTN, polyclonal, Abcam, ab14041, 1:1000; anti-SERPINE1, polyclonal, SIGMA, HPA050039, 1:50; anti-ACTA2, clone 1A4, Dako, pre-diluted) for 20 minutes (except for pan-cytokeratin, which was incubated for 30 minutes). Endogenous peroxidase activity was blocked using the Envision FLEX Peroxidase-Blocking reagent (Dako). EnVision FLEX HRP detection reagent (Dako) for secondary amplification and enzymatic conjugation. Chromogenic visualisation was performed using haematoxylin counterstaining and 2×5-minute washes in either diaminobenzidine (DAB, for pan-cytokeratin staining) Following staining for cytokeratin, sections were sequentially stained for fibroblast markers and visualised using 3-amino-9-athylcarbazole (AEC). Antigen retrieval was performed between each staining iteration to remove the previous round’s antibodies, along with removal of the labile AEC staining using organic solvents (50% ethanol, 2 minutes; 100% ethanol, 2 minutes; 100% xylene, 2 minutes; 100% ethanol, 2 minutes; 50% ethanol, 2 minutes).

#### Digital pathology processing and image analysis

Stained slides were scanned at 20x with the ZEISS Axio Scan.Z1, using ZEN 2 software (ZEISS). Image processing was carried out using custom macro scripts in the Fiji software package^48^. Images from each staining iteration were registered using the Linear Stack Alignment with SIFT plugin^49^. Then converted to “pseudo-immunofluorescence” (pIF) multi-layered TIFF images with the colour deconvolution plugin^50^, using the “H-AEC” vector matrix followed by subtracting channel 3 (“green”) from channel 2 (“red”) to distinguish between brown (Dab) staining and red (AEC) staining. Cells were then segmented based on haematoxylin (nuclear) staining and cytoplasmic regions were simulated as ellipses grown from the segmented nuclei. The mean staining intensity and fraction of cellular area positive for each stain was then calculated per cell. This, plus XY coordinates for each cell were exported for histo-cytometry analysis, using the spatstat package in R^51^.

#### Fibroblast identification and subpopulation classification

Stromal cells were identified using a random forest classifier. To optimise this classifier a subset of the histo-cytometry dataset was generated to include 2000 cells from 3 samples, which equally covered the quantiles of stromal marker expression (ensuring both stromal and non-stromal cells were equally represented). The coordinates associated with these cells were then used to manually annotate these cells as fibroblasts or stromal cells and non-fibroblasts (**Extended Data Fig. 3b**). The manually annotated dataset was then split into training (n=1500) and test (n=500) sets. Random forest models were trained to detect annotated stromal cells based on either nuclear morphology, marker expression, or both in combination (**Fig. 3b**). In the test dataset, the latter approach yielded the highest accuracy (AUC = 0.92; overall accuracy = 84.4%). This classifier was then applied to the entire histo-cytometry dataset, identifying 1.9×10^6^ stromal cells.

A random forest classifier was also used to assign stromal cells identified by histo-cytometry to the relevant subpopulation. To train this classifier, imputation was performed on the “Hub cells” dataset for *ACTA2, SERPINE1* and *POSTN* using the Seurat *AddSmoothedScore* function. Imputed expression values were then scaled and randomly split into training (66% of cells) and test (33% of cells) sets. A random forest classifier was then trained and tested on these 2 datasets respectively, demonstrating 97.3% accuracy in the test set.

Histo-cytometry estimates of cellular expression levels were calculated as Intensity scaled pixel counts. The positive pixels per 1000 pixels (PPC) and integrated density was calculated for each segmented cell. To determine pixel intensity independent of the fraction of cellular area with positive staining the Integrated density measurements were normalised for PPC (using a linear model) and scaled from 1-2, such that 1 indicates minimum intensity and 2 indicates maximum intensity. Intensity scaled pixel counts were generated as the product of this scaled intensity measurement and PPC per cell. Log2(Intensity scaled pixel counts + 1) were used to estimate the expression of each protein per cell. These expression values were then scaled to have the same range as scRNA-Seq measurements for each gene. The random forest classifier was applied to this scaled data, calculating the probability of each cell being a specific fibroblast subpopulation. In cases where the classifier failed to accurately identify the appropriate subpopulation (probability < 0.5) these cells were designated as undetermined. After applying this filter, 71.8% of stromal cells were assigned to one of the nine subpopulations.

#### Spatial Analysis

Tissue regions (Tumour, margin/edge, adjacent normal and vessels) were annotated by a consultant pathologist (GJT). Subpopulation abundance in each region was calculated using the *owin* function in the spatstat R package^51^.

DBSCAN spatial clustering was carried out using the dbscan package in R^52^. The histo-cytometry dataset was subset to include stromal cells from each sample. The DBSCAN was then used to identify spatially discrete regions of high stromal density, using a randomly down-sampled subset of each sample including 10% of all stromal cells. The minimum points parameter was set to the default of 4, as recommended for two-dimensional data. The ε (eps) parameter was set to 500 (equivalent to 110 µm), based on manual identification of the inflection point in kNN-distance plots for each sample. The predict function was then used to apply this clustering to all fibroblasts in each sample. The fraction of subpopulations at each niche was calculated and niches with similar compositions were identified by unsupervised hierarchical clustering using Ward’s method.

### Cell culture

Primary lung fibroblasts were isolated from disaggregated tissue samples as described previously^14^ and cultured in DMEM supplemented with L-glutamine (1% *v/v*; Sigma), penicillin-streptomycin (1% *v/v*; Sigma) and foetal calf serum (FCS; 10% *v/v* for routine maintenance and 1% *v/v* for experimentation; Biosera). Human Recombinant TGF-β1 (R&D Systems) was reconstituted in 4mM HCl, 0.1% (*w/v*) BSA and administered to cells at a dose of 2ng/ml.

Human recombinant IL-1β (ThermoFisher) was reconstituted in deionised water and administered at 10 ng/ml.

### Quantitative Real-time PCR (QPCR)

Cell pellets from trypsinised cells underwent RNA extraction with DNase digestion using Reliaprep™RNA Cell Miniprep System (Promega). RNA quantitation was performed using a NanoDrop Spectrophotometer (Thermo Fisher Scientific). One microgram of RNA was reverse transcribed in a 20μl reaction volume, using the High Capacity cDNA Reverse Transcription Kit (Applied Biosystems), according to manufacturer’s instructions. Two nanograms of cDNA was analysed by PCR using TaqMan Real-Time PCR Assays (COL1A1 [Hs00164004_m1]. IL6 [Hs00174131_m1]; and housekeeping genes - ACTB [Hs01060665_g1], B2M [Hs00187842_m1], GAPDH [Hs02786624_g1]; Thermo Fisher Scientific) and the QuantStudio 7 Flex Real-Time PCR system (Thermo Fisher Scientific).

### Statistical analysis

Statistical analysis was performed as described in the methods above and in figure legends, as appropriate for each statistical method implemented.

### Data Availability

The scRNA-sequencing data generated in this study will be made available using the NCBI Gene Expression Omnibus database when accepted for publication.

## Figures

**Extended Data Figure 1:**
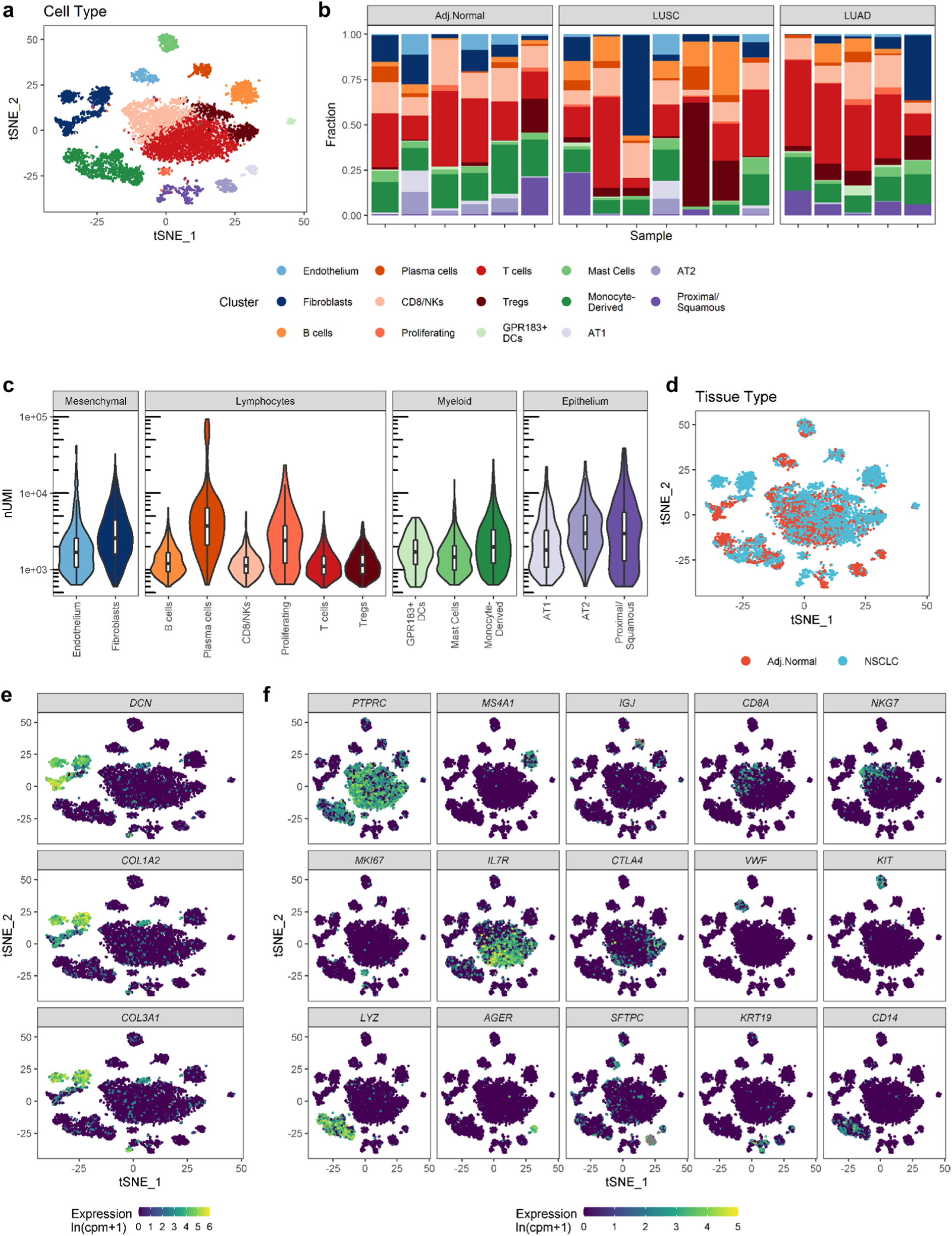
a) Human tumour-adjacent lung (n=6) and NSCLC (n=12) samples were disaggregated and analysed by scRNA-seq. A 2D visualisation (tSNE dimensionality reduction) of cell-type clustering is shown, highlighting different cell populations. b) Barplots showing the cell-type composition of each sample analysed by scRNA-seq. c) Violin and boxplots showing the number of unique molecular identifiers (UMI) per cell for different cell-types. d) tSNE plot showing the tissue type for cells analysed by scRNA-seq e) Feature plots showing the expression of fibroblast marker genes. f) Feature plots showing the expression of canonical markers for different cell-types.

**Extended Data Figure 2:**
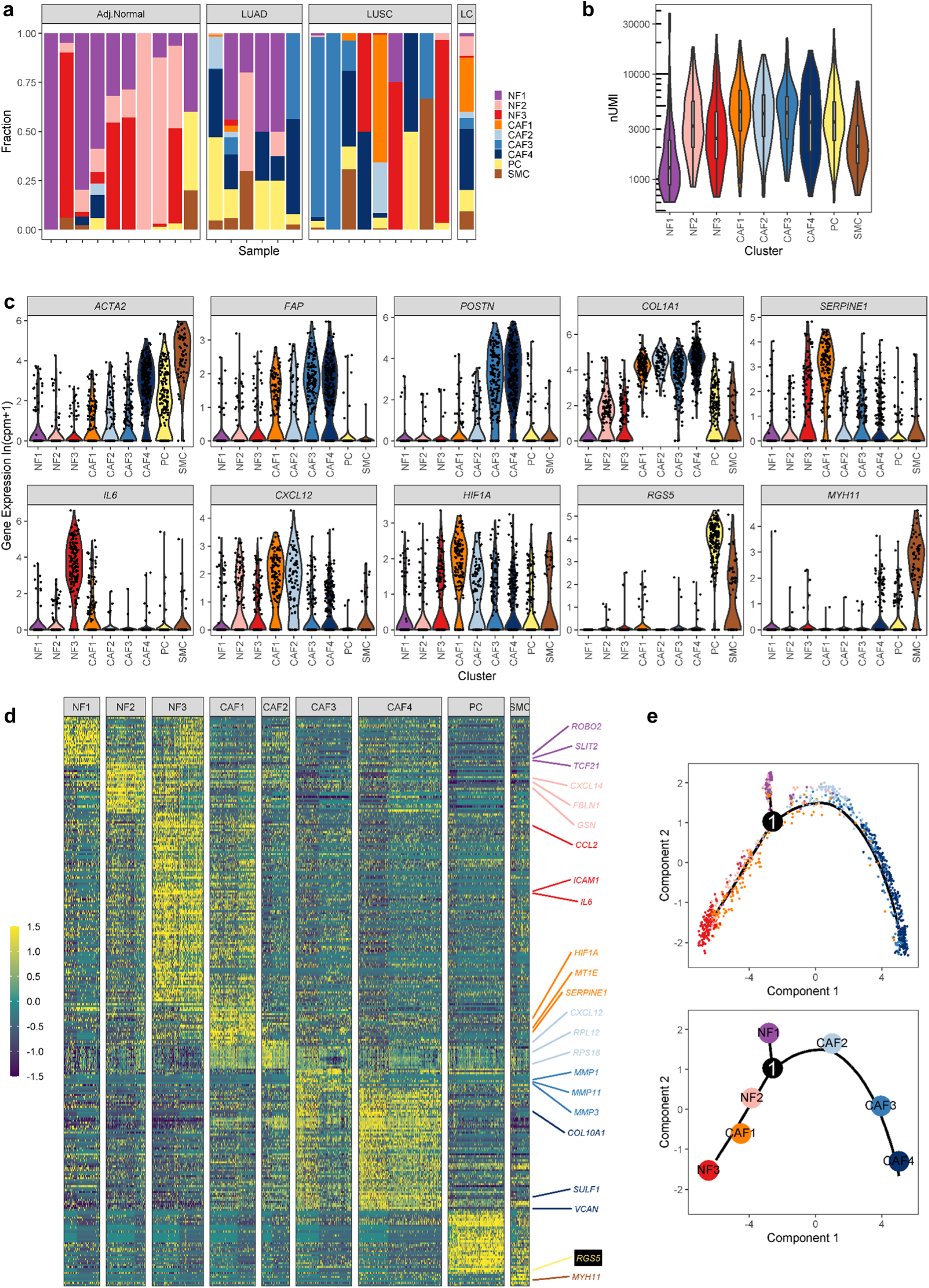
a) Barplots showing the stromal subpopulation composition of each sample analysed. b) Violin and boxplots showing the number of unique molecular identifiers (UMI) per cell for each stromal subpopulation. c) Violin plots showing the expression of selected marker genes across stromal subpopulations. d) Heatmap showing consensus cluster marker expression. Three selected markers for each fibroblast subpopulation are labelled. e) Plots showing trajectory inference validation using the Monocle algorithm

**Extended Data Figure 3:**
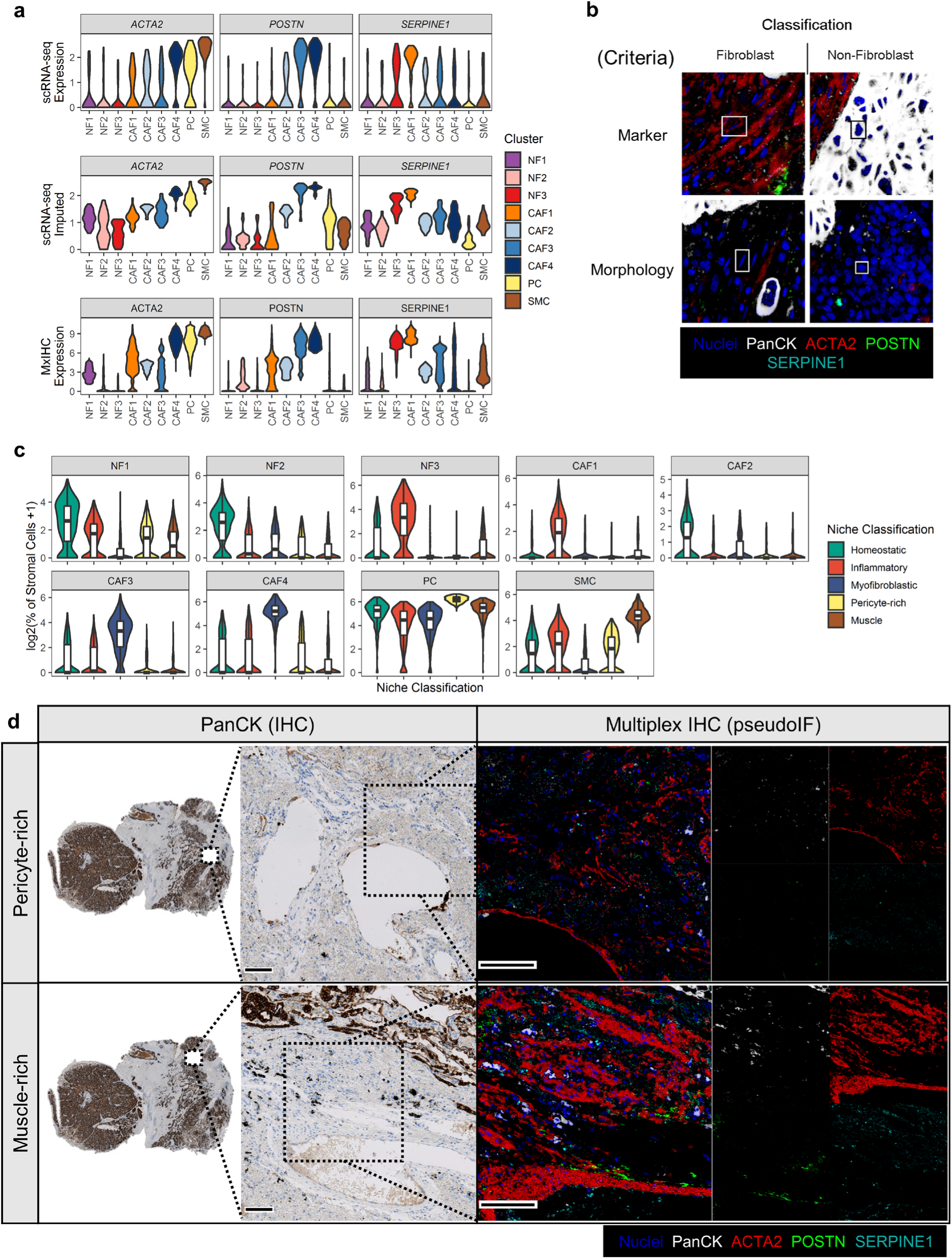
a) Violin plots showing expression of ACTA2, POSTN and SERPINE1 in the scRNA-seq and MxIHC datasets, as indicated. b) Representative images showing manual annotation of fibroblasts for training a random forest classifier for automated detection. c) Box and violin plots showing the relative fraction of stromal subpopulations across different niches. d) Representative micrographs showing a pericyte-rich and muscle-rich niche. Scale bars represent 100µm.

**Extended Data Figure 4:**
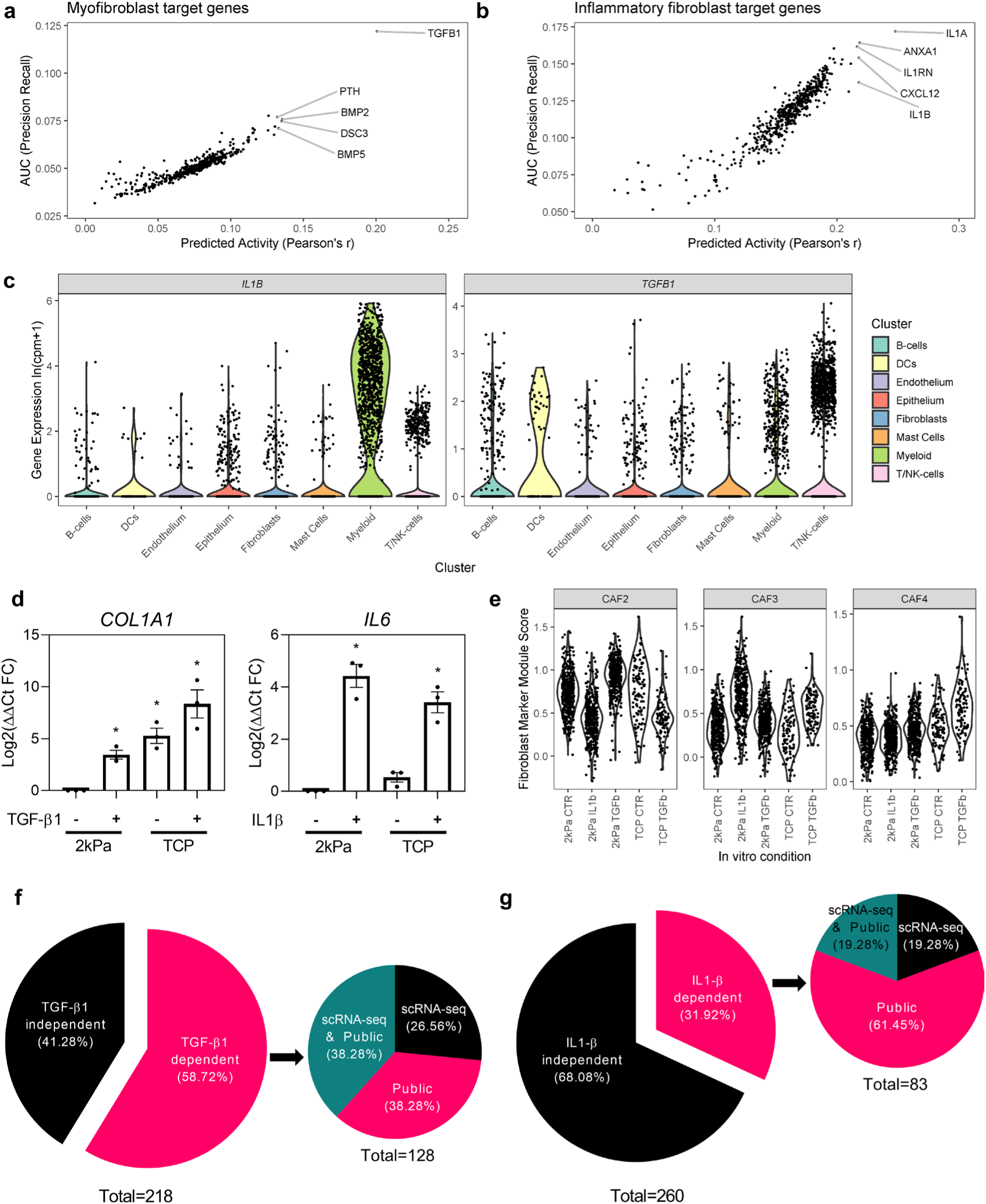
a) Scatter plots showing ligands predicted regulatory activity over myofibroblast marker or pseudotime module genes. b) As per (a) for inflammatory fibroblast marker genes. c) Violin plots showing the source of *IL1B* and *TGFB1* expression in the NSCLC tumour microenvironment, measured using the TLDS scRNA-seq dataset. d) Barplots showing QPCR analysis of primary fibroblasts (n = 3 independent experiments) cultured on a 2kPa substrate or tissue culture plastic (TCP) treated with TGF-β1 (2ng/ml for 72 hours) or IL-1β (10ng/ml for 72 hours). Statistical significance was assessed using one-way ANOVA with Dunnet’s comparison between treatment groups and the 2kPa CTR group. *adj. *p*<0.05. e) Violin plots showing relative expression (measured by scRNA-seq) of proto-myofibroblast and myofibroblast marker genes in primary fibroblasts treated as described above. f) Pie chart showing the proportion of myofibroblast genes up-regulated following TGF-β1 treatment and the relative proportion of these genes identified from our scRNA-seq analysis or publicly available datasets (see also Supplementary table 6). g) Pie chart showing the proportion of inflammatory fibroblast genes up-regulated following IL-1β treatment and the relative proportion of these genes identified from our scRNA-seq analysis or publicly available datasets (see also Supplementary table 6).

**Extended Data Figure 5:**
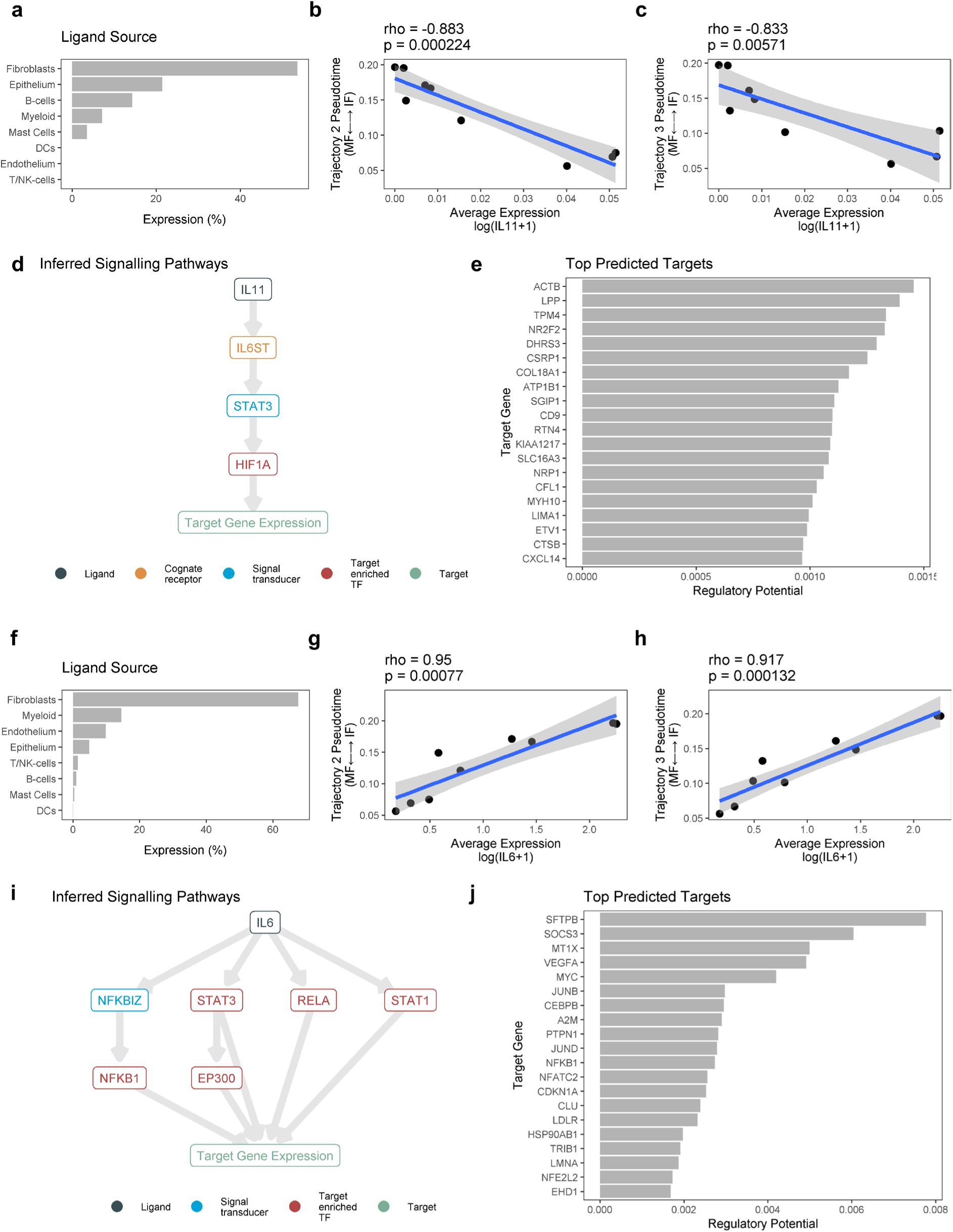
a-e) Example of signalling pathway inference for IL-11 mediated regulation of TGF-β1 independent myofibroblast genes, generated using an R shiny app available at <u>myofibroblast activation</u>. f-j) As per a-e, showing an example of signalling pathway inference for IL-6 mediated regulation of IL-1β independent inflammatory genes, generated using an R shiny app available at <u>inflammatory fibroblast activation</u>. a&f) Barplot showing the relative contribution of different cell-types to the expression of *IL11* in myofibroblast and proto-myofibroblast rich samples (a); or *IL6* expression in inflammatory and proto-inflammatory rich samples (f). b-c & g-h) Scatter plots showing the relationship between *IL11* expression and trajectory 2 (CAF4-CAF3-CAF2-NF2-NF3; b&g) or trajectory 3 (CAF4-CAF3-CAF2-CAF1-NF3; c&h) pseudotime. d&i) Inferred downstream mediators involved in IL11 mediated regulation of TGF-β1 independent myofibroblast genes (d) or IL6 mediated regulation of IL-1β independent inflammatory genes (i), identified using the NicheNet database. e&j) Barplot showing IL11’s regulatory potential score for the top 20 TGF-β1 independent myofibroblast genes (e) or IL6’s regulatory potential score for the top 20 IL-1β independent inflammatory genes (j), measured using the NicheNet database.

## Supplementary Table Legends

***Supplementary Table 1: Fibroblast subpopulation consensus marker genes***.

Differentially expressed genes were identified for each fibroblast subpopulation using a MAST test. This analysis was run on TLDS and Lambrechts datasets separately. Meta-log(fold-change) and Meta-adj.*p* values were then calculated from the lowest fold change or highest *p* value.

***Supplementary Table 2: Fibroblast subpopulation consensus marker enrichr analysis***.

Results from examining fibroblast subpopulation consensus markers for enrichment with genes from gene ontology (GO) biological processes.

***Supplementary Table 3: Fibroblast subpopulation gene signature expression***.

The expression of previously described fibroblast gene signatures was analysed across subpopulations identified by scRNA-seq. Statistical significance was assessed using a Wilcox test with fdr correction to compare the median expression level per sample for each cluster to all other clusters. The gene sets analysed are also provided in a separate sheet.

***Supplementary Table 4: Pseudotime gene expression module statistics***.

Statistics for pseudotime gene modules, identified by correlation network analysis of expression levels in trajectory-inferred pseudotime.

***Supplementary Table 5: Pseudotime gene expression module enrichr results***.

Results from examining pseudotime gene expression modules for enrichment with genes from pathways (KEGG); ligand perturbation gene sets (Ligand); microRNA targets (miRNA); and transcription factor targets (TF).

***Supplementary Table 6: Identification of myofibroblast or inflammatory fibroblast genes regulated by TGF-β1 or IL-1β respectively*** Curated gene sets for TGF-β1 and IL-1β response genes in fibroblasts. Data is taken from scRNA-seq performed in this study and publicly available datasets (provided in separate sheets).

***Supplementary Table 7: Generalised linear modelling (GLM) of ligand associations with pseudotime trajectories*** The predictive value of each ligand’s average expression for the sample’s position in pseudotime was assessed using GLM. The statistics from this analysis for each pseudotime-inferred trajectory are shown.

***Supplementary Table 8: Modelling ligand regulatory potential on either TGF-β1 or IL-1β independent myofibroblast or inflammatory fibroblast activation respectively***.

This table contains statistics from GLM and predicted regulatory potential for all ligands significantly associated with myofibroblast or inflammatory fibroblast activation (*p* < 0.05). Regulatory potential was calculated using the NicheNet R package.

## Notes

https://cjhanley.shinyapps.io/inflammatoryfibroblast_activation/

### Competing Interest Statement

The authors have declared no competing interest.

